# Decoding protein language models: insights from embedding space analysis

**DOI:** 10.1101/2024.06.21.600139

**Authors:** Pia Francesca Rissom, Paulo Yanez Sarmiento, Jordan Safer, Connor W. Coley, Bernhard Y. Renard, Henrike O. Heyne, Sumaiya Iqbal

## Abstract

Foundation models, which encode patterns in large, high-dimensional data as embeddings, show promise in many machine learning related applications in molecular biology. Embeddings learned by the models provide informative features for downstream prediction tasks, however, the information captured by the model is often not interpretable. One approach to understanding the captured information is through the analysis of their learned embeddings, which in molecular biology so far has mainly focused on visualizing individual embedding spaces. This study introduces a quantitative framework for cross-space comparison, enabling intuitive exploration and comparison of embedding spaces in molecular biology. The framework emphasizes analyzing the distribution of known biological information within embedding space neighborhoods and provides insights into relationships between multiple embedding spaces. Comparison techniques include global pairwise distance measurements as well as local nearest neighbor analyses. By applying our framework to embeddings from protein language models, we demonstrate how embedding space analysis can serve as a valuable pre-filtering step for task-specific supervised machine learning applications and for the recognition of differential patterns in data encoded within and across different embedding spaces. To support a wide usability, we provide a Python library that implements all analysis methods, available at https://github.com/broadinstitute/EmmaEmb.

## Introduction

With the growing availability of data and computing power, foundation models have become increasingly relevant in molecular biology for encoding the high-dimensional data generated by laboratory experiments. Foundation models are trained to identify the inherent relationships within the training data, uncovering complex patterns. By using large datasets as input, foundation models place individual data points into a broader context, generating representations that are informative for a range of applications (22; 10).

Protein language models (PLMs) are a specific type of foundation model. These models are based on the architectures of Large Language Models, which have revolutionized text processing in Natural Language Processing. PLMs, such as ESM2 (22) and ProtT5 (10), are trained on large amounts of protein sequence data from various species and generate representations that capture the evolutionary relationships of proteins. The learned representations, also referred to as embeddings, can be leveraged for a variety of downstream tasks, including protein structure prediction (22), function prediction (20), and variant effect prediction (7).

As more foundation models, such as PLMs, based on varying architectures, training data, and data modalities emerge, it becomes increasingly important and interesting to understand what they have learned. This understanding can guide the appropriate application of the models but also help to detect biases in the models’ representations (8). Further, understanding the models’ learned patterns enables researchers to leverage them to offer new biological insights (4). However, due to their complex architecture and high number of parameters, understanding what these models have learned is not straightforward. Therefore, it is crucial to develop methods for understanding the underlying patterns and features captured by the models.

In Natural Language Processing, embedding space analysis is a key approach for interpreting model behavior. Getting a better understanding of the distribution of points within one embedding space has helped to reveal semantic relationships (33) and biases (6) in the embeddings of language models. Additionally, identifying similarities between embedding spaces from different languages has facilitated the transfer of information from high-resource languages to a low-resource language, which has limited training data (13).

In the field of molecular biology, embedding space analysis has also been leveraged, for example, to understand the properties of the embedding space and its impact on continuous prediction tasks (1; 12), as well as to analyze multiple modalities within a single embedding space (3). However, comparisons across embedding spaces have primarily been performed visually, often using dimensionality reduction techniques such as principal component analysis, t-distributed stochastic neighbor embedding, uniform manifold approximation and projection, pairwise controlled manifold approximation, and multidimensional scaling (27; 22) or by the use of tree-based hierarchical visualizations (35). We see a potential for advancing the use of embedding space analysis in the field of molecular biology through quantitative analysis across embedding space.

A common standard for understanding which features, such as the function of a protein, are captured by a model is to benchmark the model on downstream tasks using supervised machine learning (ML) (16). Small task-specific classifiers are trained on PLM embeddings. This approach requires task-specific models for each feature and each PLM of interest, which becomes increasingly resource-intensive as the number of models and benchmark datasets grows. Analysis of the embedding space provides a less resource-intensive initial evaluation. By examining the distribution of categorical features across various embedding spaces, we can gain insights into how these features are represented by assessing the proximity of samples from different classes. Although this analysis is less detailed and does not uncover complex relationships between the embedding dimensions, it offers valuable initial insights into the informativeness of the representations across different embedding spaces and contexts. This initial evaluation is particularly useful for determining which models warrant the application of more resource-intensive ML approaches.

Additionally, by directly comparing the distances between samples in different embedding spaces, we gain insights into similarities and differences between the models’ learned information. Discrepancies in distances between the spaces can provide insights into complementary information that the models have acquired or highlight potential biases or noise the models have learned. One bottleneck hindering the cross-space embedding analysis of foundation models in molecular biology is the lack of frameworks and tools to facilitate rapid exploration. Although approaches for comparing multiple embedding spaces exist in other domains, particularly language (see Table 1), their value in molecular biology applications remains limited, and to our knowledge, the listed tools have not been applied in this context. Two key factors contribute to this gap. First, there has been a great emphasis on comparing embedding spaces visually after dimension reduction methods are applied, with little exploration of quantifying feature distributions. Second, there is often a focus on analyzing individual data points, a practice common in tools designed for Natural Language Processing, where entities such as words are easily interpretable by users. However, in molecular biology, context is essential, as the names of proteins and genes by themselves may not provide substantial meaning, even to experts. Instead, valuable insights are often gained by understanding the distribution of groups of data points with a specific biological meaning. Therefore, we emphasize the importance of incorporating the relationship between different natural groupings of the samples into the analysis, which is commonly lacking in Natural Language Processing tools.

**Table 1.**
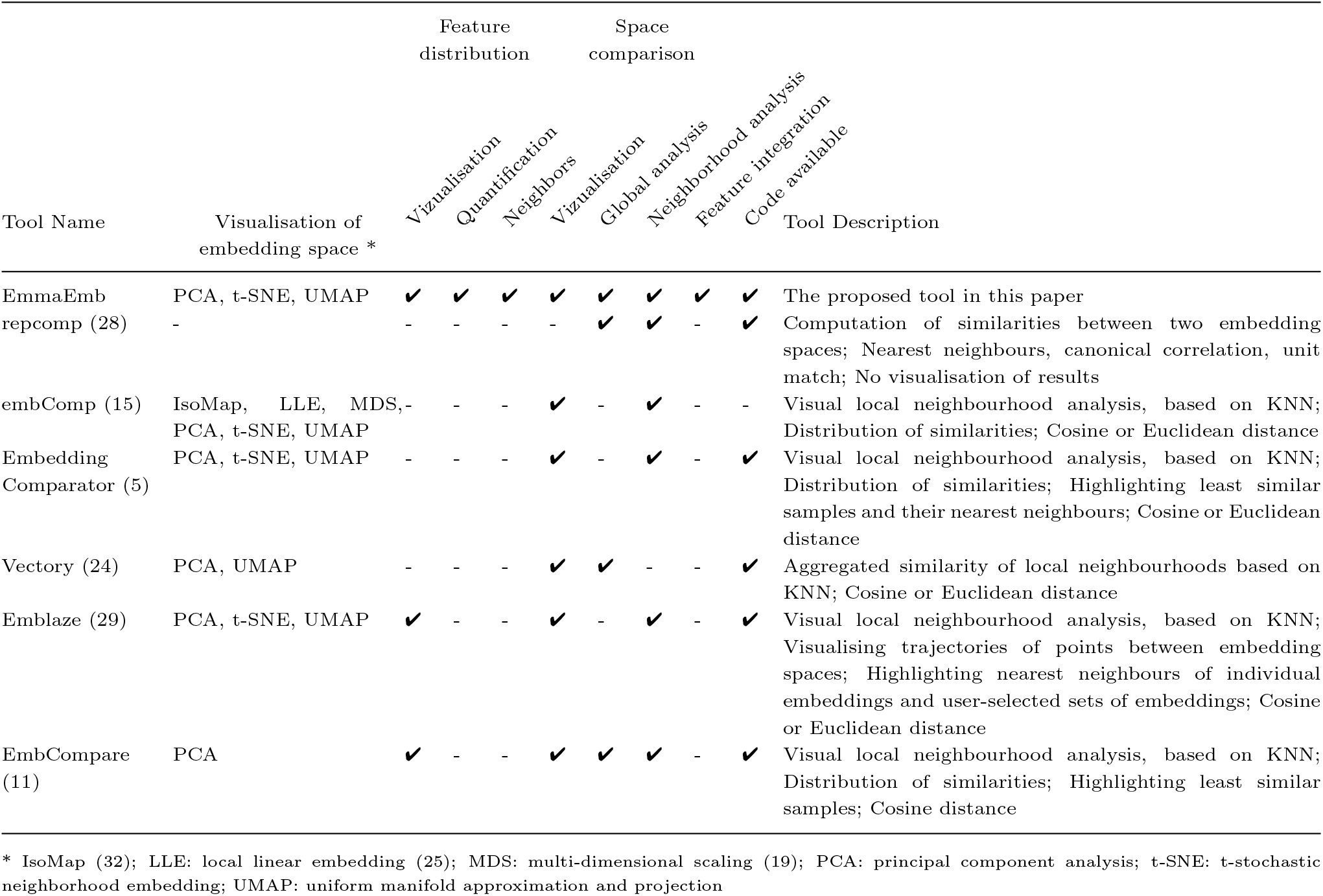
Comparison of published tools for the simultaneous analysis of multiple embedding spaces.

To bridge this gap, we developed a comprehensive framework for the quantitative and cross-space analysis of embedding spaces in molecular biology. Our approach focuses on feature distributions within embedding space neighborhoods and enables comparisons across embedding spaces using methods such as global distance metrics and local neighborhood analyses. We evaluate insights gained from our analysis framework by comparing them to those derived from task-specific ML models. Additionally, we investigate how embedding space analysis can be leveraged to generate new hypotheses about the similarities between data representations across different models. To promote accessibility, we publish the Python library <u>E</u>mbedding, <u>M</u>etadata, and <u>M</u>ultimodel <u>A</u>nalyzer for <u>E</u>xploration in <u>M</u>olecular <u>B</u>iology (EmmaEmb), which implements all framework components.

## Methods and Materials

### 1. Embedding space analysis framework

Figure 1 provides an overview of our analysis framework. We consider a set of samples 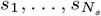, e.g. proteins, genes, etc., and *N*_*V*_ ≥ 2 embedding spaces 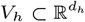, *h* = 1, …, *N*_*V*_. Every embedding space *V*_*h*_ corresponds to a different foundation model which embeds the input samples into numerical vectors, i.e., 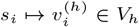, ∀ *h, i*. To analyze the information captured in the embedding spaces, a table of categorical feature vectors 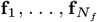, where each is a vector of length *N*_*s*_ containing the feature values of each sample. By **f**_*p*_[*i*] we denote the value of the *p*-th feature of *s*_*i*_. The feature data can for example include biological natural groupings, such as protein family information, or experimental data, such as functional annotations.

**Fig. 1.**
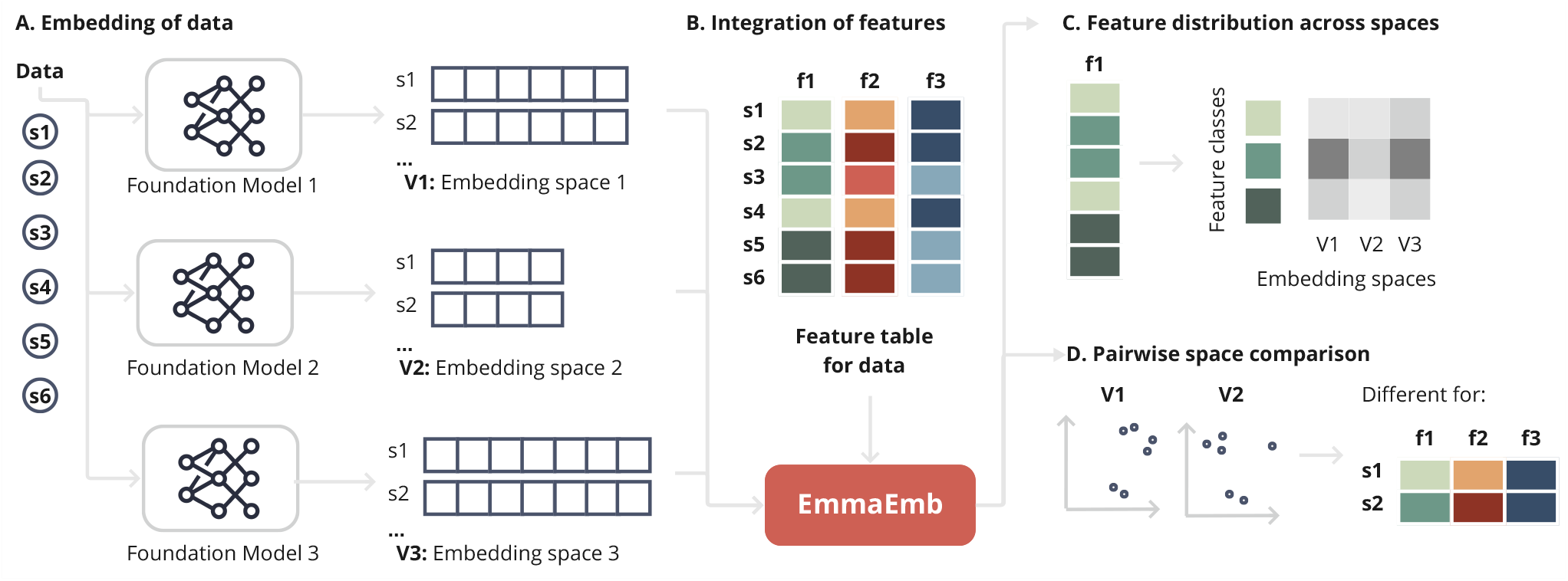
Overview of the proposed analysis workflow. Our framework aims to analyse the information captured in different embedding spaces. **A**. Starting with a set of samples (e.g. genes or proteins, *here*: Data **s**_**1**_ − **s**_**6**_), embeddings are derived from different foundation models (*here*: Foundation Models 1-3). In the example of proteins, embeddings could be derived from protein language models that vary in their architecture or training. **B** Categorical data about the samples (*here*: **f**_**1**_ − **f**_**3**_) is integrated into the analysis, e.g. annotations of the function of proteins, protein families, etc. The embeddings, along with sample-specific categorical feature data, are then analyzed together using two main types of approaches. First, **C**. deriving the distribution of specific features, such as protein families, within neighborhoods of each embedding space, and comparing them across embedding spaces. This analysis is based on a *k*-nearest neighbors approach, quantifying how closely features from one class are in proximity and which classes are neighboring classes. Secondly, **D**. two embedding spaces can be compared to identify similarities and differences in the pairwise distances between embeddings or *k*-nearest neighbors. Regions of high difference can be further analyzed using the feature data to characterize the samples that were represented differently by the model. The introduced EmmaEmb python framework is a freely accessible implementation of the described approach.

The embedding spaces may vary in dimensions, and there is no direct correspondence between dimensions across embedding spaces. To facilitate comparison across models, we focus our analysis on the distances between embeddings within each space. For each pair of samples *s*_*i*_ and *s*_*j*_, the distance between their respective embeddings, dist 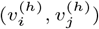, is measured in each embedding space *V*_*h*_. The pairwise distance measures form the foundation for the following analysis. Different distance measures, such as Euclidean, cosine, and Manhattan distance, can and should be used to inspect the proximity of data points from varying viewpoints (21). For parts of the analysis, only the *k*-nearest neighbors (KNN) are considered to focus on local parts of the embedding space. KNN based metrics have been tailored for various applications (34), here we adjust and leverage them for the needs of embedding comparisons. The neighbors are identified based on pairwise distance measures, with users defining the distance metric and the number of neighbors *k*. We suggest setting *k* to around 10% of the total number of samples, but it will not be larger than the size of the smallest feature class.

#### 1.1 Analysis methods

We gathered a set of quantitative methods that are suitable to measure information across embedding spaces. We group the methods into feature distribution and pairwise space comparison methods (see Figure 2). The choice of analysis steps depends on the research goals, and we do not suggest that all methods must be applied or follow a particular sequence. Each method provides a unique perspective on what the foundation models have learned, enabling insights tailored to specific use cases.

**Fig. 2.**
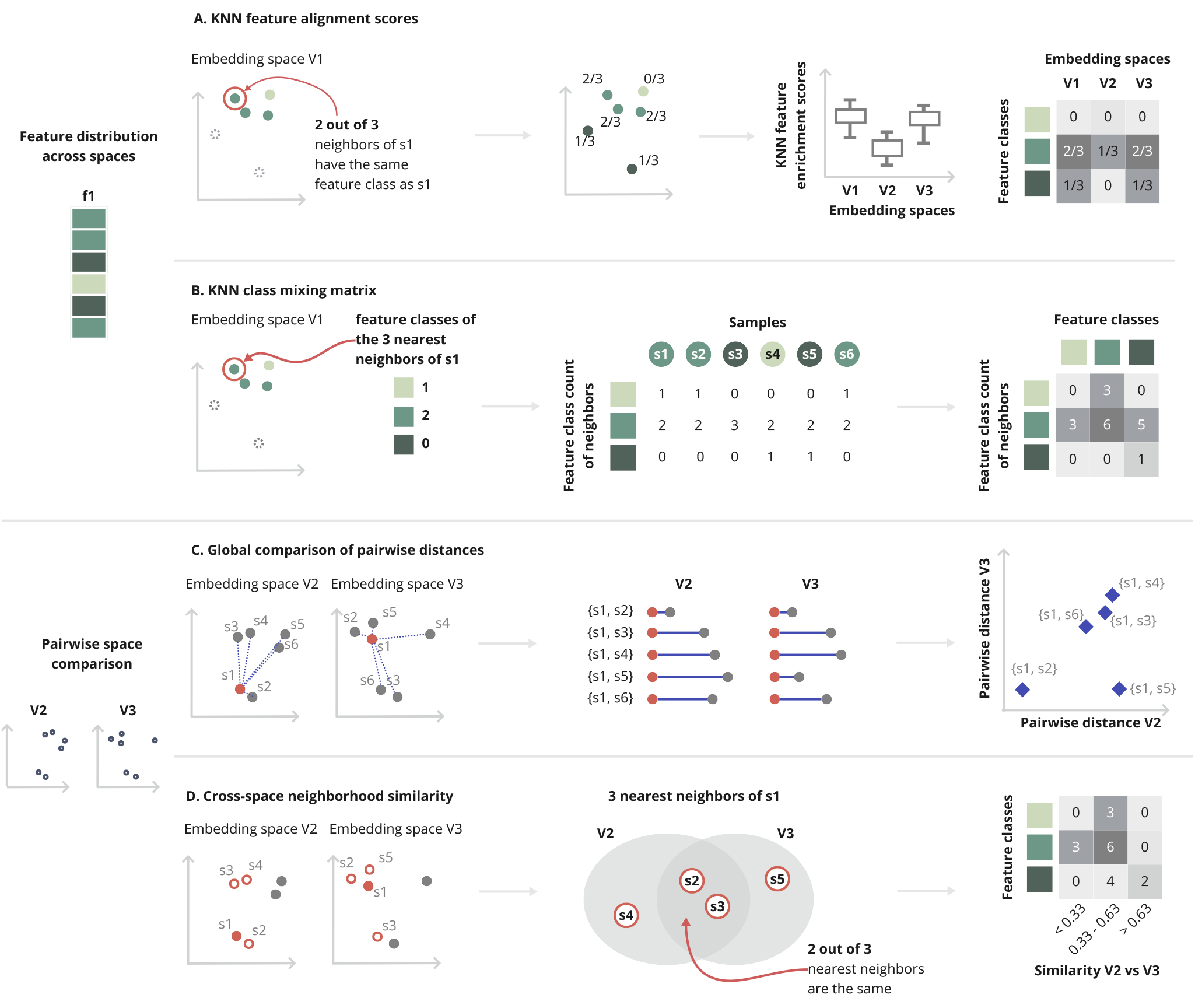
Graphical overview of proposed methods for embedding space analysis. For the analysis of the feature distribution of a specific feature (*here* : feature **f**_**1**_) for a set of samples (*here*: *s*_1_ − *s*_6_) across embedding spaces (*here*: feature **V**_**1**_ − **V**_**3**_) we propose two methods. Firstly, **A. KNN feature alignment scores**. The KNN feature alignment scores are calculated for each embedding within its embedding space, as a fraction of its *k*-nearest neighboring embeddings that share the same class label as the embedding itself. By comparing the distribution of KNN feature alignment scores across different embedding spaces, we can assess how closely embeddings of the same **f**_**1**_ class are to other embeddings of the same class. The score provides insights into how well an embedding model has learned to group embeddings with similar characteristics. Secondly, **B. KNN class mixing matrix**. The matrix is calculated separately for each embedding space (*here*: for *V*_1_). For each embedding, the class distribution of class labels of its *k*-nearest neighbors is computed. The class distributions are then summed for all embeddings of the same class. The resulting matrix shows, for each class, the number of *k*-nearest neighbors of embeddings from this class for another class. This allows insights into which classes are close to each other and which are more differentiated within the embedding space. We also propose methods for pairwise embedding space comparison, such as **C. Global comparison of pairwise distances** (*here*: between embedding space *V*_2_ and *V*_3_). For this method, we first calculate all pairwise distances between embeddings within each embedding space. Then, we compare the correlation of pairwise distances, e.g. distance between *s*_1_, *s*_2_, across the two embedding spaces. This method provides an overview of the global similarity between embedding spaces and reveals groups of samples that are differently represented. **D. Cross-space neighborhood similarity**, in contrast, quantifies local differences between two embedding spaces. It is calculated for each sample as the number of *k*-nearest neighbors shared between *V*_2_ and *V*_3_. Identifying and characterizing samples with high divergence in their local neighborhood can help to uncover structural differences between the embedding spaces.

##### 1.1.1 Feature distribution across spaces

###### KNN feature alignment scores

We define scores that provide insight into how closely related a sample is to its neighbors in terms of the feature **f**_*p*_.

For a given feature **f**_*p*_ and embedding space *V*_*h*_, the KNN feature alignment score for each sample is defined as the fraction of KNN 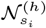 that share the same class label as the sample.

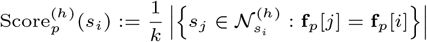

The score ranges from 0 to 1, with higher values indicating a greater representation of the feature class in the sample’s neighborhood.

KNN feature alignment scores can be stratified by feature labels to examine the extent to which each feature class is captured in an embedding space. Furthermore, KNN feature alignment scores of one embedding space can be directly compared to those from another embedding space, enabling an assessment of how well different models capture relationships for the same feature.

###### KNN class mixing matrix

We propose the inspection of how close individual feature classes are in each of the embedding spaces to gain insights into feature classes how similar different feature classes are encoded by the models. For each sample in a given embedding space and for a selected feature, we count how many times each feature class occurs within the KNN embeddings. For each feature class, we then sum the number of times the neighbors belong to that class for all samples that share the same class label. This process is repeated for every feature class, aggregating the class counts across all samples in the embedding space to get a comprehensive view of how feature classes are distributed within the space. The results are presented in a matrix format, where each column corresponds to a specific feature class, and each row represents the count of neighbors for samples labeled with that feature class, organized by their own feature class. High values in the matrix suggest that embeddings from the feature class in the column are frequently found close to samples from the feature class in the row, indicating that the model interprets them as more similar. However, it is important to note that a high degree of neighborhood coherence for a feature class may also imply the presence of multiple subclusters within that class.

##### 1.1.2 Pairwise space comparison

###### Global comparison of pairwise distances

To perform a global comparison of pairwise distances across embeddings, we compute Spearman’s rank correlation coefficient of pairwise distance values. This statistical measure allows us to evaluate how well the pairwise relationships between embeddings are preserved across different embedding spaces by comparing their ranking across different spaces.

To visualize, the pairwise distances in the two embedding spaces can be plotted against each other in a scatter plot.

###### Cross-space neighborhood similarity

To quantify and analyze local differences between two embedding spaces, we first identify the KNN for each sample in both spaces. As in previous approaches for embedding space comparison (15; 5; 29; 11), we compute the overlap of these KNN between the two spaces for each sample, enabling a comparison of how similar the local neighborhoods of each sample are in the two embedding spaces. We focus on the most and least similar sets of samples. In addition to previous work, we quantify the feature distribution within these subsets to characterise the differing samples and provide context.

#### 1.2 Implementation in the EmmaEmb Python library

To make the proposed framework accessible, we developed a light-weight modular Python library called EmmaEmb. The library is centered around the `Emma` object, which handles embedding spaces, feature data, and intermediate computational results. Multiple functions on the `Emma` object enable analysis of feature distributions across embedding spaces, including feature alignment and neighborhood analysis, as well as pairwise space comparisons between different embedding spaces. EmmaEmb integrates functionalities from established libraries such as scikit-learn (23) and Plotly (17) to calculate the described analysis measures and offers intuitive options for visualizing the results. For distance measurement, EmmaEmb provides a variety of metrics, including Manhattan, Euclidean, and cosine distances, along with their normalized and scaled variants. The library is designed as a modular framework that can be extended for further types of analysis, such as clustering approaches or topological analysis. Additionally, the repository includes a script that facilitates the extraction of embeddings from the PLMs listed in Table 2, integrating sequence-chopping algorithms to process long sequences.

**Table 2.**
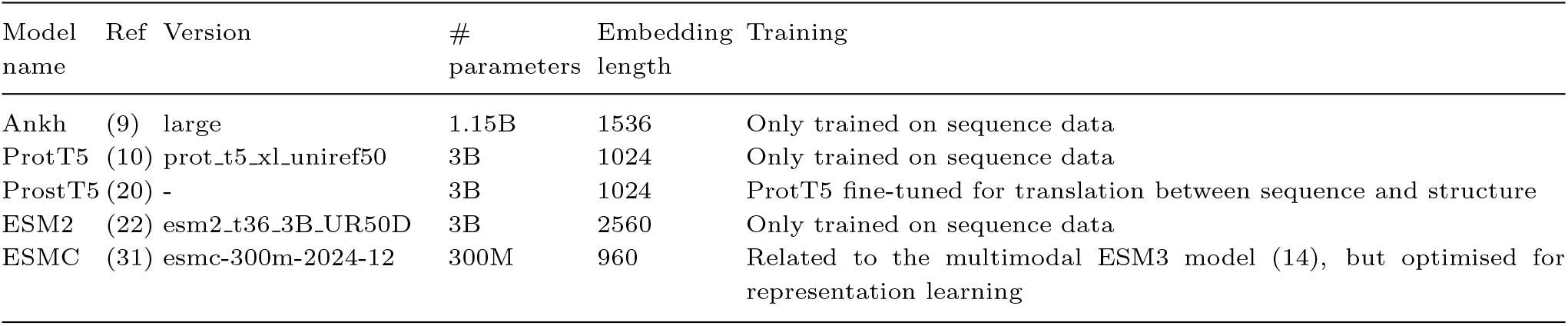
Overview of the protein language models used in this study.

### 2. Experimental setup

#### 2.1 Datasets

Two datasets were chosen to test the capabilities of our embedding space analysis framework.

##### DeepLoc

We utilized the training set of version 1.0 of the DeepLoc dataset, which consists of 11,231 protein sequences from eukaryotes annotated with subcellular localization information across 10 distinct classes (2). DeepLoc is a widely used benchmark dataset for evaluating protein subcellular localization prediction models (30; 16), offering a comprehensive resource for comparing the embedding space analysis framework’s results to task-specific supervised ML model performance.

##### PLA2G2

The PLA2G2 dataset consists of protein sequences from the phospholipase A2 family, specifically the phospholipase A2 Isozyme Group II (PLA2G2) published in (18). The dataset includes protein sequences annotated across 7 different species and proteins are categorized into 11 different enzyme classes. To enable comprehensive data integration, only proteins with both species and enzyme class information were included, resulting in a final set of 446 proteins. Compared to the DeepLoc dataset, the PLA2G2 dataset exhibits greater sequence similarity, making it more suitable for studying evolutionary changes.

#### 2.2 Embeddings

Table 2 provides an overview of the transformer-based protein language models evaluated in the experiments. As some of these models have restrictions on the maximum sequence length, protein sequences longer than 1022 amino acids were divided into smaller chunks. Each chunk was processed with a minimum overlap of 300 amino acids to maintain sequence continuity. The embeddings for each chunk were then aggregated using the mean of all chunk embeddings, resulting in a single unified representation for the full-length protein. As proposed by Brandes et al. (7), to ensure that the entire sequence is effectively represented, chunks were taken with overlapping windows from both sides, and if necessary, an additional window was included in the middle to fill any gaps.

## Results and discussion

### Distribution of protein localization information

We evaluate our embedding space analysis approach by comparing its results with those derived from training ML models on embeddings, focusing on the subcellular localization task. In particular, we consider results from light-attention networks, which have demonstrated superior performance compared to other approaches on embeddings from PLMs for this task (30). The networks were trained specifically for the subcellular localization task using the DeepLoc training set (30; 16), and we compare our findings to the results of these task-specific models reported on test sets. KNN computation in the embedding space analysis was conducted using *k* = 100 and the cosine distance.

To compare how well models capture the subcellular localization information we calculated the KNN feature alignment scores. According to the estimated distribution, ProtT5 has the highest scores on average, followed by Ankh, ESM2 and ProstT5 (Figure 3A). This ranking was statistically validated using one-sided Wilcoxon signed-rank tests (n = 11,231): ProtT5 vs ESM2 (p-value *<* 0.0001, ranked biserial correlation coefficient = 0.20), ESM2 vs Ankh (p-value *<* 0.0001, ranked biserial correlation coefficient = 0.12), and Ankh vs ProtT5 (p-value *<* 0.0001, ranked biserial correlation coefficient = 0.46). This ranking aligns with previously reported performance when the embeddings from these models were used with a trained light-attention network for this task (16). The observed consistency between the mean KNN feature alignment scores and ML-based performance rankings shows that embedding-based KNN feature alignment analysis can serve as a proxy for evaluating the informativeness of embeddings in downstream tasks.

**Fig. 3.**
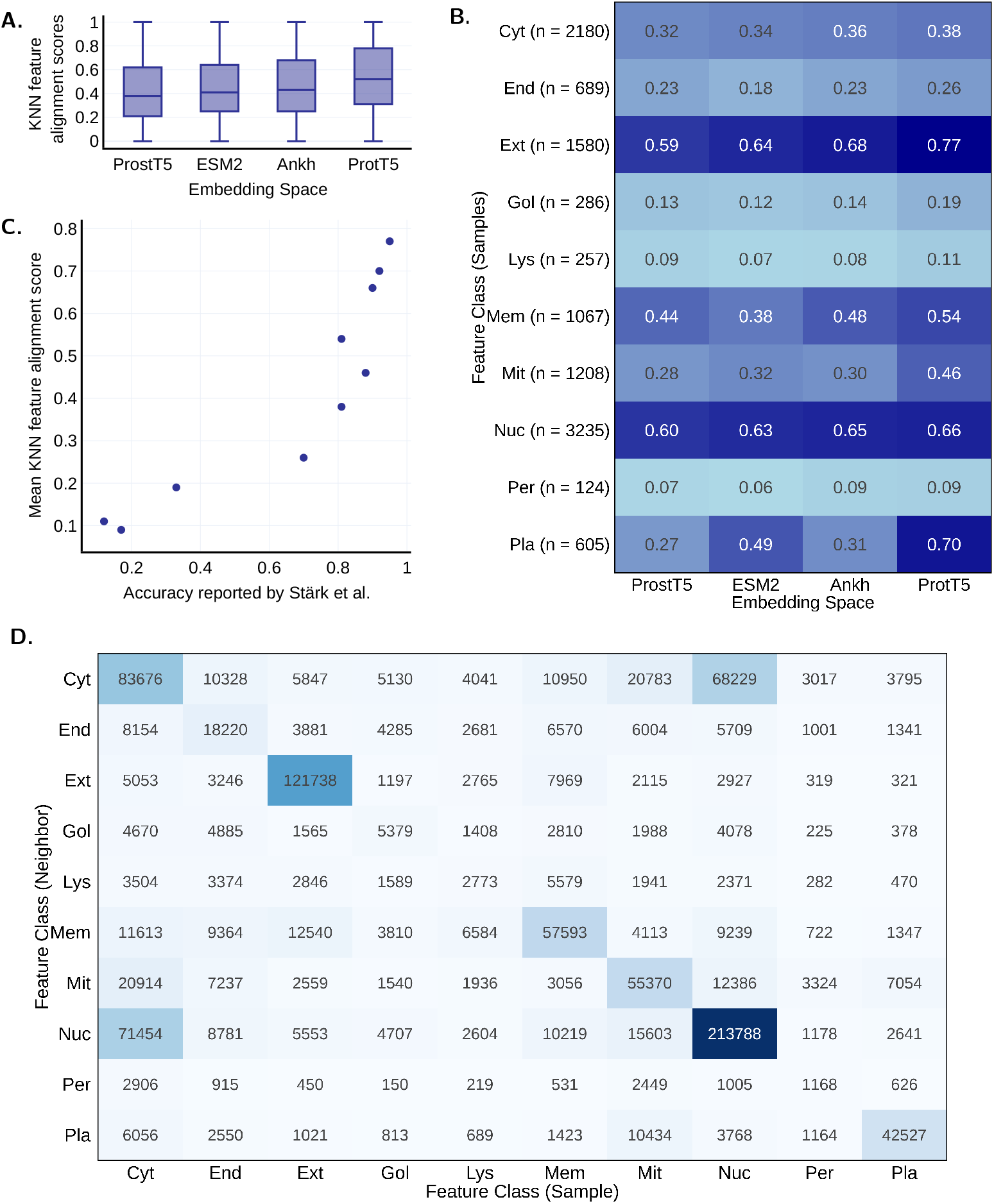
Feature distribution analysis of subcellular localization feature in the DeepLoc dataset across the ProstT5, ESM2, Ankh, and ProtT5 models. KNN analysis is performed with k = 100 and cosine distance. **A**. Boxplots displaying the KNN feature distribution scores per model. Protein localization information is most closely represented in ProtT5, followed by ESM2, Ankh, and ProstT5. **B**. Heatmap of the mean KNN feature distribution score per embedding space and feature class. Some classes are strongly represented across all models, while others are captured by only specific models. **C**. Scatter plot comparing the accuracy of a supervised machine learning model trained on the ProtT5 embeddings (as reported by Stärk et al. (30)) on the x-axis and the mean KNN feature alignment scores for the ProtT5 embeddings on the y-axis. Each dot represents a different localization class, and a strong correlation is observed between these two measures. **D**. KNN class mixing matrix for the ProtT5 embedding space. Some classes are represented in closer proximity to each other than others. Cyt: Cytoplasm; End: Endoplasmic reticulum; Ext: Extracellular; Gol: Golgi apparatus; Lys: Lysosome/vacuole; Mem: Cell membrane; Mit: Mitochondrion; Nuc: Nucleus; Per: Peroxisome; Pla: Plastid.

Next, we calculated mean aggregated KNN enrichment scores per embedding space, stratified by feature classes (Figure 3B). This approach highlights which feature classes are consistently well captured in the model’s representations, such as “Extracellular”and “Nucleus”, while also identifying models that capture specific classes more effectively, such as ProtT5 for “Plastid”and “Mitochondrion”. Further analysis reveals a strong correlation between the mean KNN feature alignment scores of the ProtT5 embedding space for specific subcellular localization classes and the prediction accuracy achieved by a light-attention model trained on ProtT5 embeddings, as reported in (30). This relationship, illustrated in Figure 3C, is supported by a two-sided Spearman’s rank correlation coefficient of 0.97 (p-value *<* 0.0001). This analysis provides similar insights but at a more granular level, demonstrating that embedding space analysis can be used to approximate the models’ overall understanding of not only of specific features but also for different feature classes.

Even more fine-grained, the differentiation of different feature classes within the ProtT5 embedding space are also examined using the proposed KNN class mixing matrix (see Figure 3D). For classes that are not yet well differentiated in the space, such as “Cytoplasm”, it is possible to dive deeper into feature classes that proteins from this class are located nearby and might be “mixed”with. For example, proteins classified as “Cytoplasm”are in proximity to those classified as “Nucleus”. This information can aid in more targeted training of models to better differentiate specific features and improve class separation in the embedding space. Again we compare the results with the results from the light-attention approach. For each class, we examine the closest neighboring class in the KNN class matrix and then identify the classes to which the proteins were most frequently misclassified in task-specific ML classification. We found that for all 8 of 10 classes, the closest neighboring class corresponded to the class where proteins were most often misclassified as belonging to that class in the task-specific ML model.

Our results support the idea that, with an increasing number of models and benchmarking tasks, embedding space analysis could play a key role in pre-selecting models for further, more computationally intensive, task-specific, and embedding-specific analyses. It is important to recognize that task-specific ML models may uncover task-relevant complex relationships between dimensions that might not be apparent in a global analysis on the entire embedding space. Adding to this, recent work comparing the performance of task-specific ML models trained on embeddings versus fine-tuning PLMs suggests that the superior performance of one PLM over another in a task-specific embedding approach does not necessarily predict its superiority after fine-tuning (26). Therefore, benchmarking for specific tasks remains essential for evaluating task-specific models.

### Comparison of ESMC and ProtT5 embeddings for PLA2G2 proteins

In this section, we apply our pairwise space comparison to the PLA2G2 dataset. We compare the ProtT5 model against the recently published ESMC model. The two models differ in their training, parameter sizes, as well as embedding dimensions (see Table 2). Our aim is to investigate whether these models exhibit differences in how they represent this set of enzyme proteins.

Global correlation analysis of the pairwise cosine distances between the embeddings of the models shows a high Spearman’s rank correlation coefficient of 0.75 (p-value *<* 0.0001) between ESMC and ProtT5. This indicates a substantial similarity in how these two models represent relationships among the sequences overall.

When examining the scatter plot of pairwise cosine distances for embeddings of the ESMC and ProtT5 models, we observe that many data points align along a linear trend, consistent with the high correlation value. However, a distinct cluster of proteins deviates profoundly, with much greater pairwise distances in ProtT5 compared to ESMC (see Figure 4A.1). We could identify this driven by a subset of bird proteins as illustrated in Figure 4A.2. Notably, this difference in representation is not observed when comparing pairwise distances for other species, such as crocodiles (see Figure 4A.3). This analysis raises new questions about the models and why they have learned to represent a subset of bird proteins differently. One possibility is that one of the models has learned finer distinctions between subgroups of bird species, potentially due to differences in training data.

**Fig. 4.**
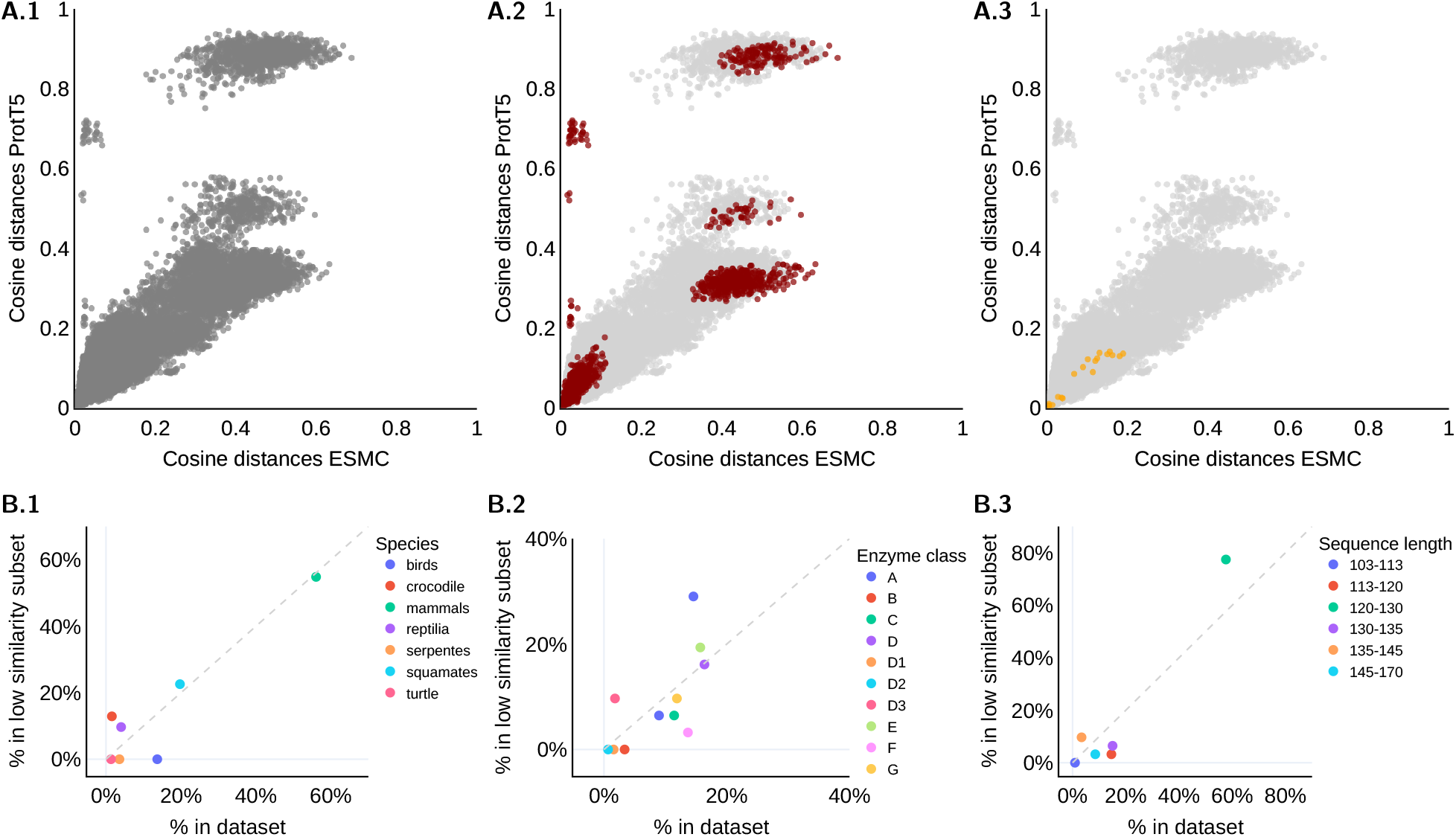
**A**. The top row displays pairwise cosine distances between PLA2G2 embeddings using ESMC (x-axis) and ProtT5 (y-axis). **A.1** for all embeddings, **A.2** between embeddings of bird proteins (red), and **A.3** between embeddings of crocodile proteins (yellow). **B**. The bottom row illustrates the class distribution of proteins with 30% local neighborhood overlap (cosine distance, *k* = 10) between the ESMC and ProtT5 models, compared to overall dataset representation (x-axis) for the features: **B.1** species, **B.2** enzyme classes, and **B.3** sequence length.

As a final step, we compare local neighborhoods between the models and identified proteins with a local neighborhood similarity of less than 30% when comparing the ESMC and ProtT5 models. Analyzing the distribution of species, enzyme classes and amino acid sequence lengths (see Figure 4B). In comparison to the global analysis, local changes between the ESMC and ProtT5 models were not observed in bird proteins but rather in proteins from squamates and reptilia. This suggests that, even though the positions of points corresponding to bird species may have shifted within the embedding space, the local neighborhood information for bird sequences is preserved, indicating a consistent representation of their relationships within the space. This analysis highlights the importance of analyzing embedding spaces at both global and local levels, which will give partially complementary views.

## Conclusion

In summary, we show that comparing embedding space analysis with supervised machine learning models trained on the embeddings provides similar insights into model learning, without requiring task-specific or model-specific training. While embedding space analysis is limited to the information captured in the overall embedding space without accounting for complex interactions or subspaces, it can play a key role in pre-selecting the growing number of models for further training.

Further, we explore the use of embedding space analysis to investigate discrepancies between PLMs on subsets of the data.

This approach helps to identify discrepancies in the information captured by the models, which can reveal gaps in the training data or highlight complementary learned information.

Although our approach is demonstrated here with protein sequence embeddings, it is domain-agnostic. It can be applied to embedding spaces across foundation models in molecular biology and beyond, particularly in cases foundation models are combined with structured metadata.

## Author contributions statement

P.F.R. conceived and designed the study with input from P.Y.S.. P.F.R. conducted the experiments and analyses. J.S. contributed to the development of the codebase used in the analysis. P.F.R. drafted the manuscript. P.Y.S., J.S., C.W.C., B.Y.R, H.O.H., S.I. provided critical feedback and input throughout the study. All author reviewed and approved the final version.

## Acknowledgments

We would like to express our gratitude to Katharina Baum, Jakub Bartoszewicz, Melania Nowicka, Marta Lemanczyk, Theresa Hradilak, Eugenia Alleva, and Hilary Finucane for their valuable discussions and insightful input regarding this project. The project was supported by the Designing for Sustainability research program, a joint initiative of the Morningside Academy for Design at MIT and the Hasso Plattner Institute, generously funded by the Hasso Plattner Foundation as well as the DFG, German Research Foundation (project number 459422098).

